# Effects of early life stress on bone homeostasis in mice and humans

**DOI:** 10.1101/2020.07.21.214122

**Authors:** Karin Wuertz-Kozak, Martin Roszkowski, Elena Cambria, Andrea Block, Gisela A. Kuhn, Thea Abele, Wolfgang Hitzl, David Drießlein, Ralph Müller, Michael A. Rapp, Isabelle M. Mansuy, Eva M. J. Peters, Pia M. Wippert

## Abstract

Bone pathology is frequent in stressed individuals. A comprehensive examination of mechanisms linking life stress, depression and disturbed bone homeostasis is missing. In this translational study, mice exposed to early life stress (MSUS) were examined on bone microarchitecture (μCT), metabolism (qPCR), and neuronal stress mediator expression (qPCR) and contrasted with a sample of depressive patients with or without early life stress by analyzing bone mineral density [BMD] (DXA) and metabolic changes in serum (osteocalcin, PINP, CTX-I). MSUS mice showed a significant decrease in NGF, NPYR1, VIPR1 and TACR1 expression, higher innervation density in bone and increased serum levels of CTX-I, suggesting a milieu in favor of catabolic bone turnover. MSUS mice had a significantly lower body weight compared to control mice, and this caused minor effects on bone micro-CT. Depressive patients with experiences of childhood neglect also showed a catabolic pattern. A significant reduction in BMD was observed in depressive patients with childhood abuse and stressful life events during childhood. Therefore, future studies on prevention and treatment strategies for both mental and bone disease should consider early life stress as a risk factor for bone pathologies.

## 1. Introduction

Bones are essential components of the musculoskeletal system and subjected to continuous remodeling as an adaptation mechanism to environmental changes. Disturbances in bone development and remodeling by formation/resorption of the extracellular matrix (ECM) or differentiation of osteoblasts into osteocytes and apoptosis of osteocytes could result in reduced bone mass and increased fracture risk.

Beside well-documented mechanisms like menopause-associated hormonal changes, aging-related factors, changes in physical activity, as well as drugs and diseases, recent research suggested that psychosocial stress and stress-linked neuronal processes are associated with an increased risk for bone pathologies [1-5]. Autonomic and sensory nerve fibers densely innervate bone, predominantly in the periosteum and in areas with high metabolic activity and bone turnover. Neuronal structures can make direct contact with osteoblasts and osteoclasts, which in turn express receptors for neuronal markers on their surface and can produce neuronal mediators in an auto- and paracrine fashion [6-10]. Importantly, sympathetic and peptidergic neuronal factors can act as stress mediators and appear to play a key role in bone turnover [11-13]. Of specific relevance are tyrosine hydroxylase (TH), a rate-limiting enzyme for noradrenaline synthesis, neuropeptide Y (NPY), often co-expressed with noradrenaline, vasoactive intestinal peptide (VIP), often co-expressed with acetylcholine, and the neuropeptides calcitonin gene related peptide (CGRP) and substance P (SP) that are expressed in sensory nerve fibers. In addition, neurotrophic growth factors such as nerve growth factor (NGF) and brain derived neurotrophic factor (BDNF) that guide outgrowing nerve fibers to bony areas requiring innervation are of interest. Stress alters their expression, and they can also act as direct and indirect growth factors for osteoblasts [14-26]. Latest studies indicate that psychosocial stress can lead to structural and functional changes in neuronal plasticity, neuronal marker expression, mitochondria and inflammation [4, 27, 28], possibly resulting in downstream alterations of bone homeostasis [29-33].

In humans, major early life stress like childhood maltreatment strongly affects the stress response, as well as behavioral and cognitive functions throughout life [34, 35]. Strong associations were shown between childhood maltreatment and higher risk for psychological and physiological disorders including depression, hypersensitivity of the neuroendocrine, autonomic and peptidergic stress response leading to altered dynamics of the hypothalamic pituitary adrenal (HPA) axis, and changes in the neuronal mediators that can influence bone health [36]. Childhood maltreatment is one of the best-documented risk factors for depression [37-40] and depression in turn has been associated with a higher risk for reduced bone mineral density (BMD), osteoporosis and bone fractures [41-45].

However, existing knowledge about underlying mechanisms between early life stress, neurogenic stress response elements, and detrimental changes in bone metabolism and microstructure/BMD has been limited by several reasons. First, although early life stress like childhood maltreatment is a frequent phenomenon, it is hardly accessible to retrospective and longitudinal studies due to its rare diagnosis and ethical requirements for case-control studies. Second, access to biological material and strict control of confounding factors, such as physical activity and eating habits, are challenging in human studies [46, 47]. Animal models offer an obvious way to investigate major stress experiences early in life despite these obstacles. Animal models are helpful counterparts of human studies to investigate underlying mechanisms in a highly controlled environment that allows extensive tissue sampling, strict definition of adverse events and confounding factors, and lower subject variability.

Unpredictable maternal separation and maternal stress during early life (MSUS paradigm [48]) is an experimental stress paradigm in rodents that can mimic aspects of childhood maltreatment and induce long-lasting health effects, such as increased depression-like behavior (anhedonia) and altered brain activity in adulthood [49]. The MSUS paradigm severely impacts physiology and behavior in the exposed animals but also in their descendants across several generations via mechanisms involving epigenetic factors in the germline [48, 50-52].

To address the knowledge gap between early life stress, stress-dependent neurogenic markers that mark neuronal plasticity and detrimental changes in bone metabolism and bone microstructure/BMD, we first examined the impact of early life stress on neuronal mediator expression (qPCR) and bone microarchitecture (μCT), metabolism (qPCR on bone, ELISA on serum for osteocalcin OC, PINP, CTX-I), and in mice using the MSUS paradigm (**Aim 1**). Then we translated the findings to humans by analyzing metabolic changes in serum (OC, PINP, CTX-I) and microarchitecture (DXA) in a sample of depressive patients with or without retrospectively reported experiences of early life stress, like childhood maltreatment and stressful life events during childhood (**Aim 2**).

## 2. Results

### 2.1. Mouse Study

#### 2.1.1. Bone innervation and neuronal mediators

We first assessed the effects of early life stress on neuronal signaling and neurogenic factors in bone metabolism by measuring neuronal density and expression patterns of neurogenic markers in the bone of MSUS and control mice (*n =* 6-8 each).

To study changes in neuronal marker gene expression, qPCR was performed. Gene expression analysis revealed that key receptors for nerve growth factors, neurotransmitters and neuropeptides (***Figure 1A***) and selected receptors ligands (***Figure 1B***) were changed in MSUS mice in a pattern that may promote higher bone turnover and lower bone density. In detail, mRNA expression of the receptors NPYR1 (p < 0.05, ***Figure 1A***), VIPR1 (p < 0.001, ***Figure 1A***) and TACR1 (p < 0.01, ***Figure 1A***) were significantly reduced in MSUS mice whole bone homogenates (after removal of bone marrow), suggesting their presence on osteocytes. These three receptors are involved in the regulation of proliferation and differentiation in bone and their concomitant downregulation may lead to higher remodelling activity with less proliferation and more differentiation of bone cells. Furthermore, a significant reduction of the neurotrophin NGF (p < 0.001, ***Figure 1B***) was revealed, while NPY showed a downtrend in MSUS (p = 0.068, ***Figure 1B***). This would hamper both neuronal plasticity and bone remodelling in the face of a challenge such as a wound or aging leading to osteoporosis. Other markers of bone innervation and metabolism such as NGFR, ADRB2, CHRNA7, RAMP1, and TAC1 were similar in control and MSUS mice, whereas TRKA, TRKB, TH, ChAT and VIP were below the detection limit.

**Figure 1A:**
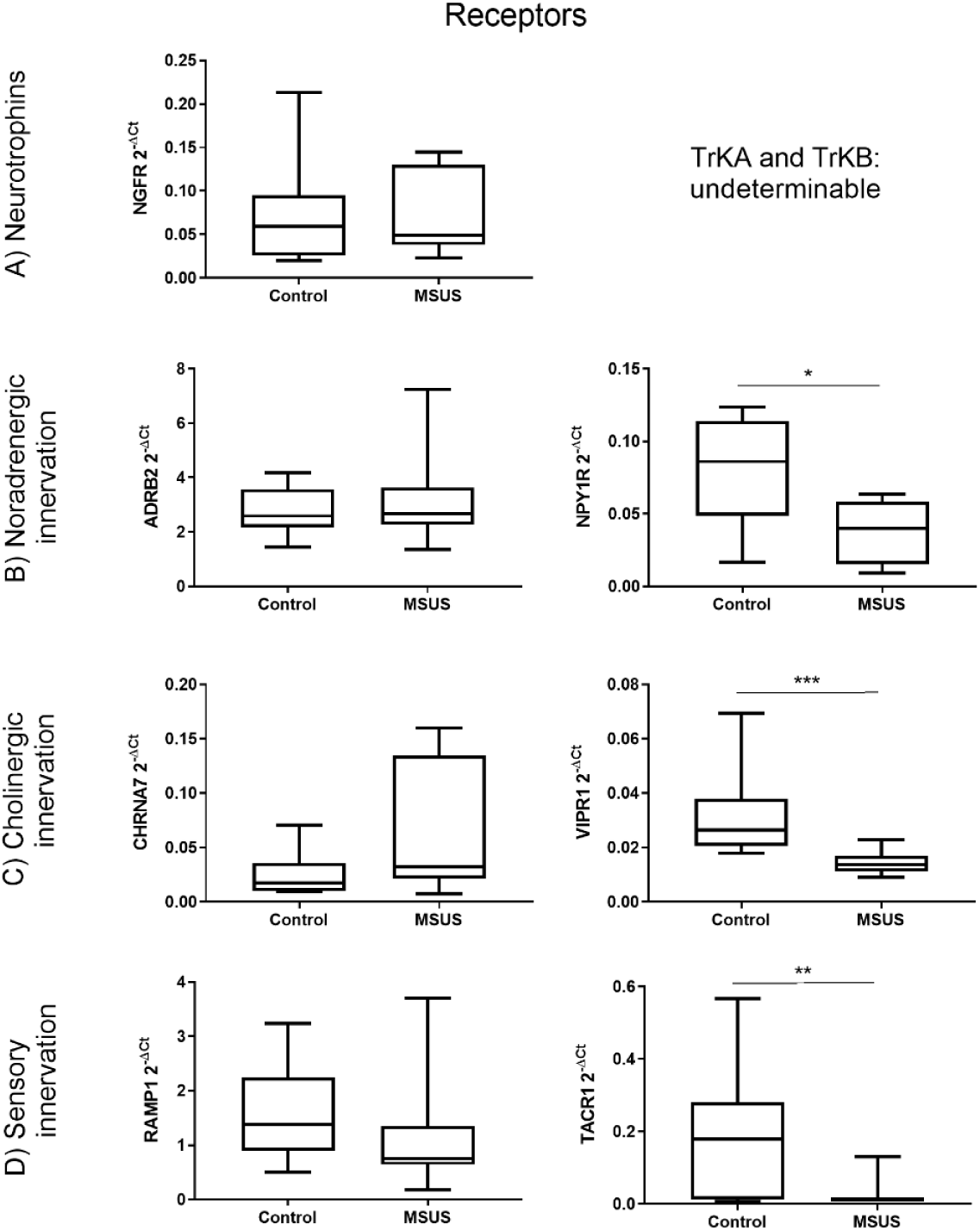
Differential gene expression of neuronal receptors in bone between control and MSUS mice. Shown are the gene expression of neurotrophin receptors **(A)**, noradrenergic receptors **(B)**, cholinergic receptors **(C)** and receptors involved in sensory innervation **(D)**. Data (n = 8 per group) are expressed as Min to Max of 2^-dCt^. Significance level: ***p < 0.001, **p < 0.01 and *p < 0.05 between indicated groups.

**Figure 1B:**
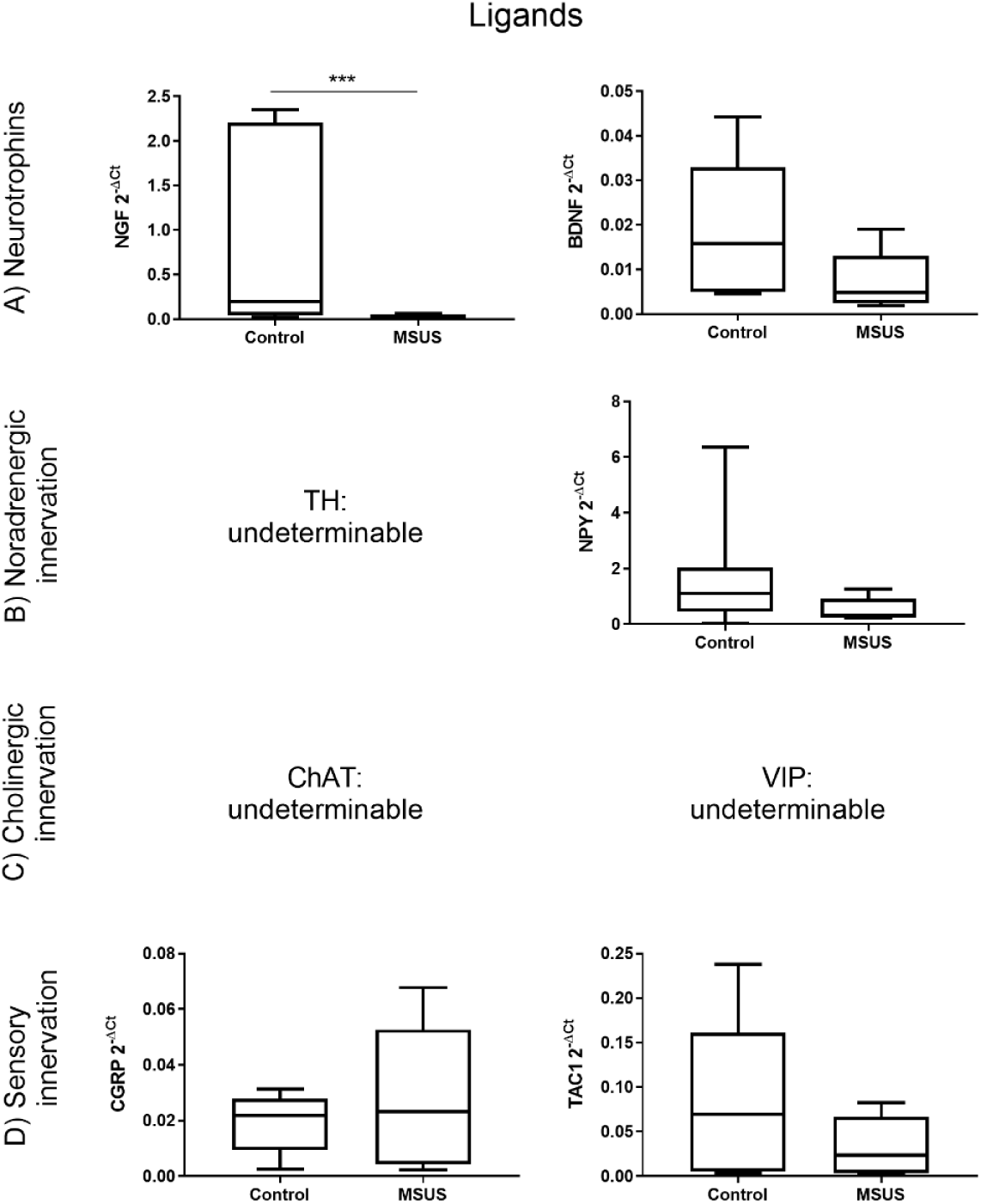
Differential gene expression of neuronal ligands in bone between control and MSUS mice. Shown are the gene expression of neurotrophins **(A)**, noradrenergic ligands **(B)**, cholinergic ligands **(C)** and ligands involved in sensory innervation **(D)**. Data (n = 8 per group) are expressed as Min to Max of 2^-dCt^. Significance level: ***p < 0.001 between indicated groups.

We subsequently investigated bone innervation by immunohistomorphometry. The density of nerve fibers labeled with the pan-neuronal marker PGP 9.5 was significantly increased in the bone of MSUS mice (***Figure 2***), whereas nerve fibers labeled with the neuronal plasticity marker GAP 43 were unaffected (not shown), indicating that higher innervation was not due to increased neuronal plasticity, but was likely a stable feature. Higher innervation density potentially leads to higher sensitivity of bone neurons to stimuli like injury. Noteworthy, the neuropeptides NPY, VIP and SP were undetectable by immunohistochemistry, most likely due to insufficient bone preservation. This likely results from the post-mortem fixation method used for ethical requirements, which does not preserve small peptides (the eleven amino-acid sequence of SP for example is rapidly digested by endogenous enzymes after termination of circulation) [53-55].

**Figure 2:**
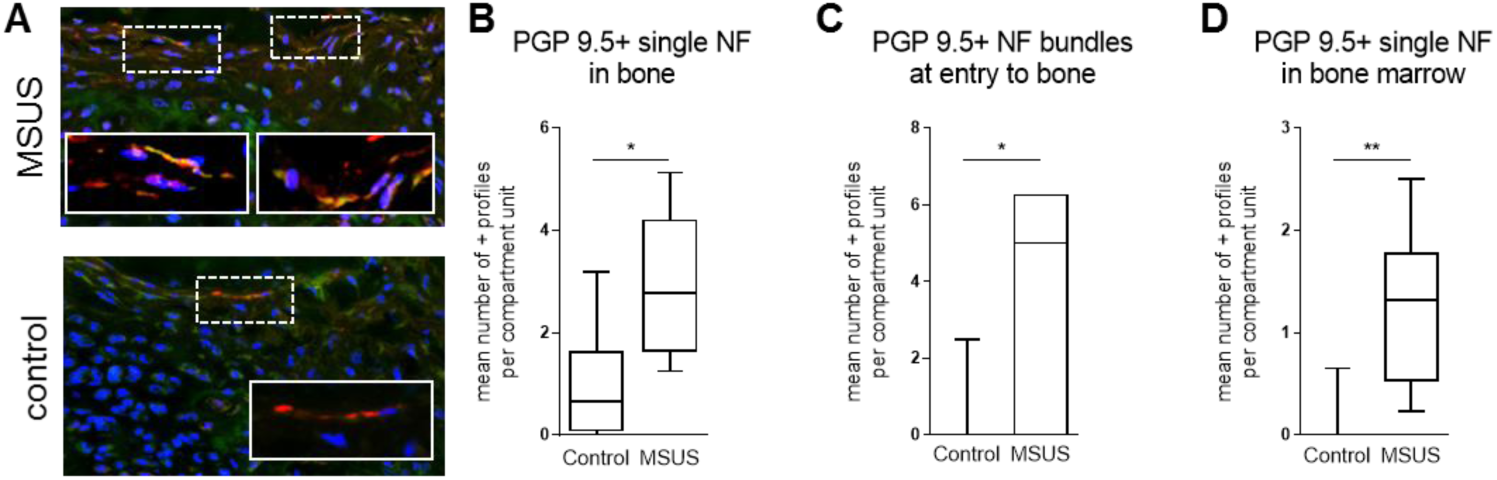
Increased bone innervation in MSUS mice. Representative images of the third tail bone from control and MSUS mice labeled with PGP 9.5 pan-neuronal marker for immunofluorescence histomorphometry of nerve fibers (NF - red) in bone. Nuclei are counterstained with DAPI (blue), mast cells with FITC-avidin (green) **(A)**. Shown are the increased mean numbers of immune-positive profiles per compartment unit of single NF in bone **(B)**, NF bundles at entry to bone **(C)** and single NF in bone marrow **(D)**. Data (n = 8 Control, n = 6-7 MSUS) are expressed as Min to Max of number of positive profiles per compartment unit. Significance level: **p < 0.01 and *p < 0.05 between indicated groups.

#### 2.1.2. Bone metabolic parameters

We then examined if changes in neuronal markers were associated with bone metabolic changes by measuring markers in serum and bone. Serum levels of osteocalcin and PINP, two markers associated with bone formation, did not change after MSUS (***Figure 3A-B***). In contrast, the level of CTX-I, which is associated with bone resorption, was significantly increased in MSUS mice compared to control mice (p < 0.01, n=14 each; ***Figure 3C***).

**Figure 3:**
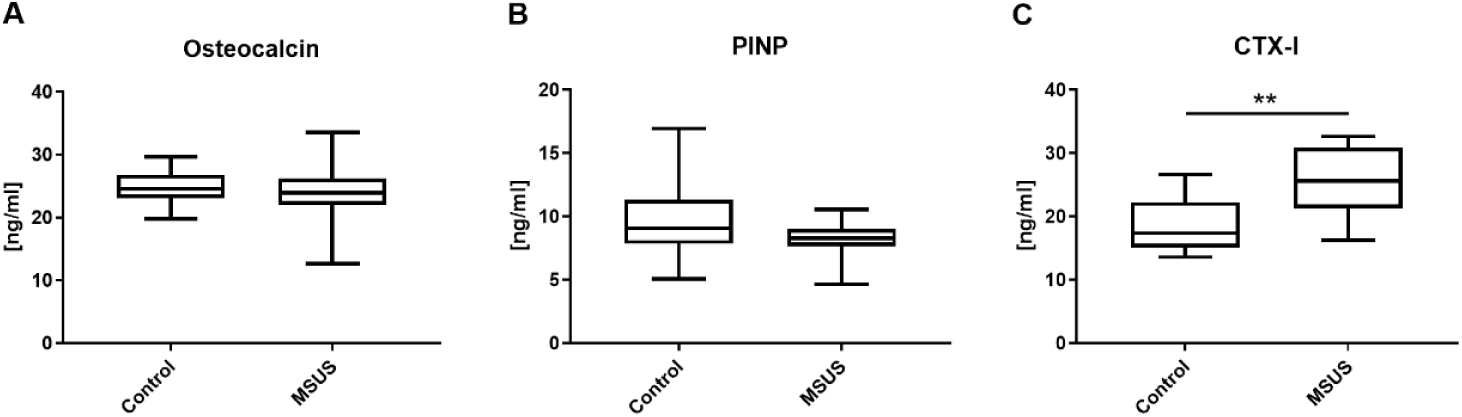
Bone formation and resorption markers in serum of mice. Shown are serum concentration measurements of osteocalcin **(A)**, procollagen type 1 N-terminal propeptide (PINP) **(B)** and c-terminal telopeptide of type I collagen (CTX-I) **(C)** in mice. Data (n=14 per group) are expressed as Min to Max of concentration in serum [ng/ml]. Significance level: **p < 0.01 between indicated groups.

However, this catabolic shift was not reflected by altered gene expression of tissue-specific extracellular matrix markers (osteocalcin, osteoprotegerin, osteopontin, sclerostin) in bone samples of MSUS (***Figure 4A-D***) (*n=8* each).

**Figure 4:**
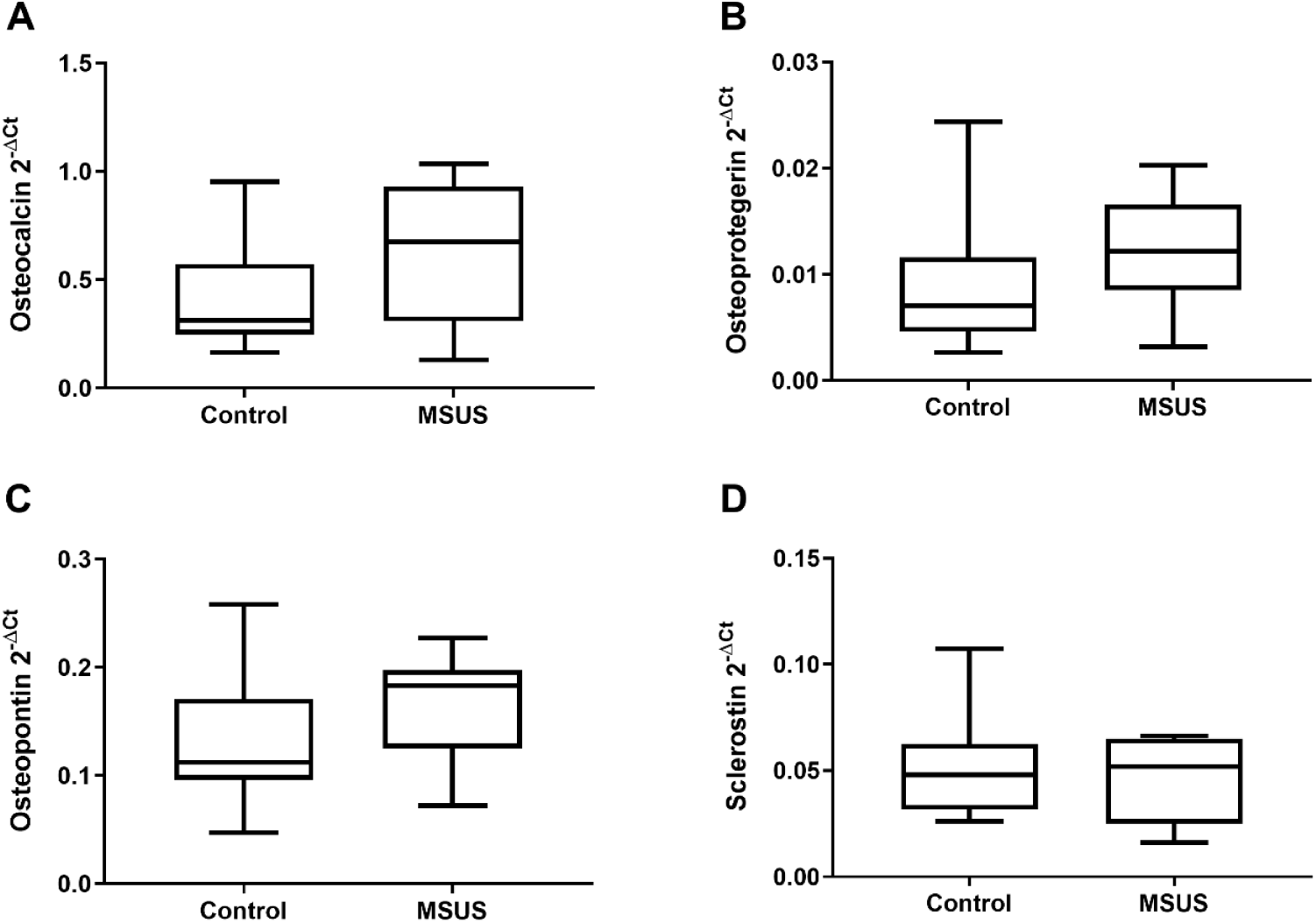
Gene expression of extracellular matrix in bone of mice. Shown are the gene expressions of tissue-specific extracellular matrix markers in bone of controls and MSUS. Genes include osteocalcin **(A)**, osteoprotegerin **(B)**, osteopontin **(C)** and sclerostin **(D)**. Data (n=8 per group) are expressed as Min to Max of 2^-dCt^. Significance level: all p > 0.0

#### 2.1.3. Bone microarchitecture

Next, it was examined if MSUS affects bone microarchitecture in mice using micro-CT (n=14 MSUS, n=13 Control). As bone microarchitecture is sensitive to mechanical loading, we also measured body weight of experimental animals over time. Body weight was significantly lower in MSUS mice compared to control mice starting at postnatal day 7 (PND7) and remained lower in adulthood (p< 0.001 for PND 7-28 and p < 0.05 for 10 months, ***Supplementary Figure 1***), and this caused minor effects on bone microarchitecture.

After adjusting micro-CT data for body weight, a comparison of means between control and MSUS mice did not reveal any difference in full, cortical or trabecular microarchitecture (all values p > 0.05) (***Supplementary Table 1***). We also used blocking in tertiles for body weight and conducted a two-factorial multivariate analysis of variance, which revealed that indeed the effects of MSUS versus controls were driven by body weight. To specify this effect, we conducted sets of linear regression analyses to test for simple mediation effects and found that body weight mediated the effect of MSUS on full length, TV, BV, MV and BS (all p’s for direct effects > 0.05).

**Table 1.**
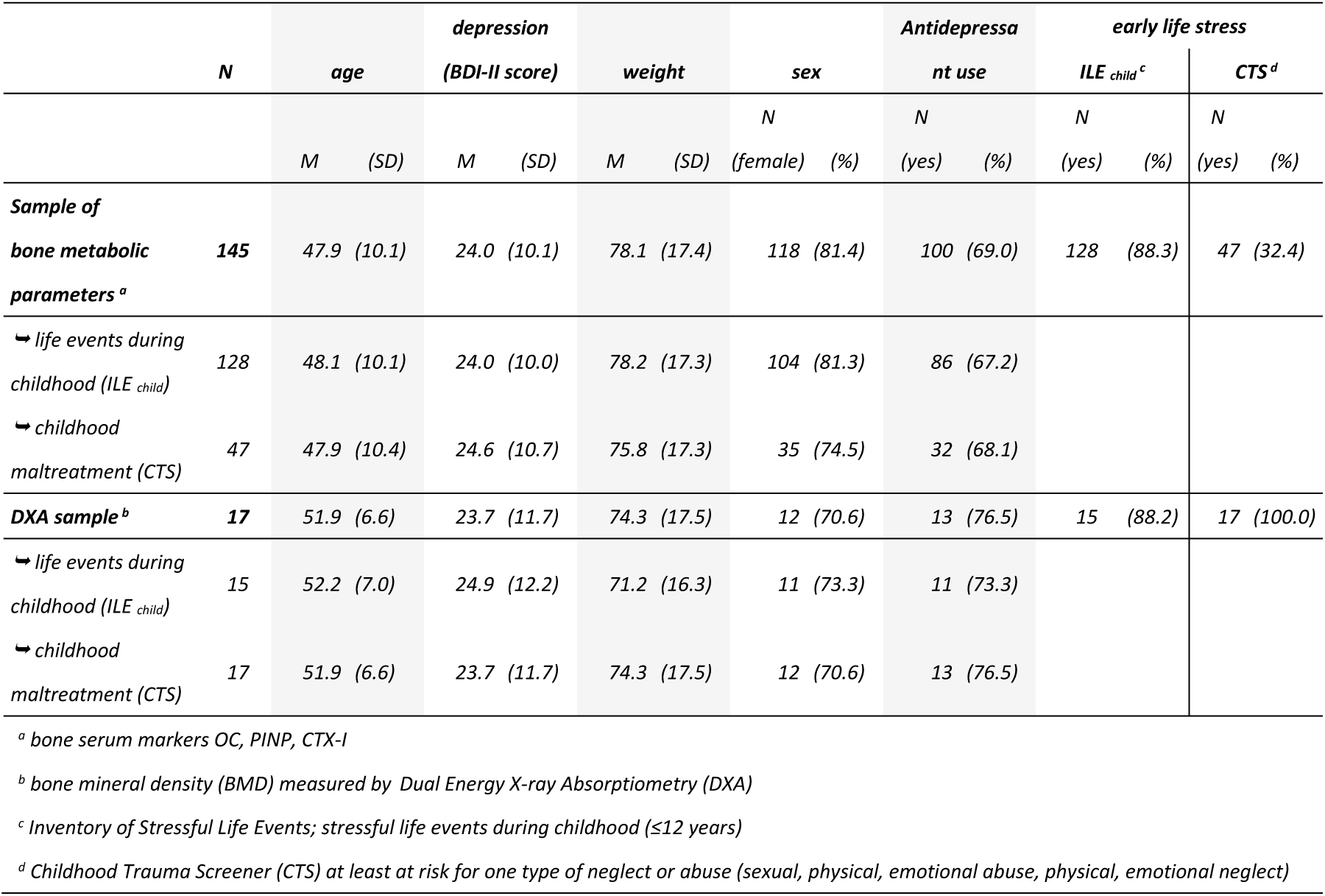
Descriptive data on sociodemographic and clinical characteristics of the human sample

### 2.2. Human Study

#### 2.2.1. Descriptive results

*N*=193 out of *n*=208 patients under study suffered from depression with minimal (35%) up to moderate/ severe (65%) severity (BDI-II: *M* = 23.8, *SD* = 10.7) and were included in further analysis. For the analysis of bone metabolic parameters, blood samples from n=145 patients were available, of whom *n* = 145 filled in the *Inventory of Stressful Life Events* (ILE) and *n=*54 the *Childhood Trauma Screener* (CTS). For the BMD analysis, n=17 patients took part in DXA bone densitometry measurements. Out of these participants, *n=*15 patients filled in the *Inventory of Stressful Life Events* (ILE) and *n=*17 the *Childhood Trauma Screener* (CTS). Missing data is due to incomplete questionnaires or drop out. Table 1 provides descriptive data on age, gender, antidepressant use, weight and early life stress for the subsamples. Average bone mineral density (BMD) in the sample was *M*= 0.942 g/cm^2^ (*SD* = 0.186 g/cm^2^) in the trochanter region of the proximal femur.

#### 2.2.2. Bone metabolic parameters

Regarding changes in human bone metabolism following early life stress, bone serum markers OC, PINP, CTX-I were analyzed. It was shown that depressed patients with childhood trauma (CTS, CTS abuse) had an increased anabolic bone metabolism in comparison to depressed patients without early life stress (CTS: *p* = 0.05* for CTX; CTS abuse: *p* = 0.03* for P1PN, ***Table 2***). This anabolic bone metabolism was likewise found for depressed patients with stressful life events during childhood (ILE _child_) in comparison to depressed patients without early life stress (*p* = 0.04* for P1PN, ***Table 2***). Depressed patients with experiences of childhood neglect (CTS neglect) lack such anabolic reaction (***Table 2***).

**Table 2.**
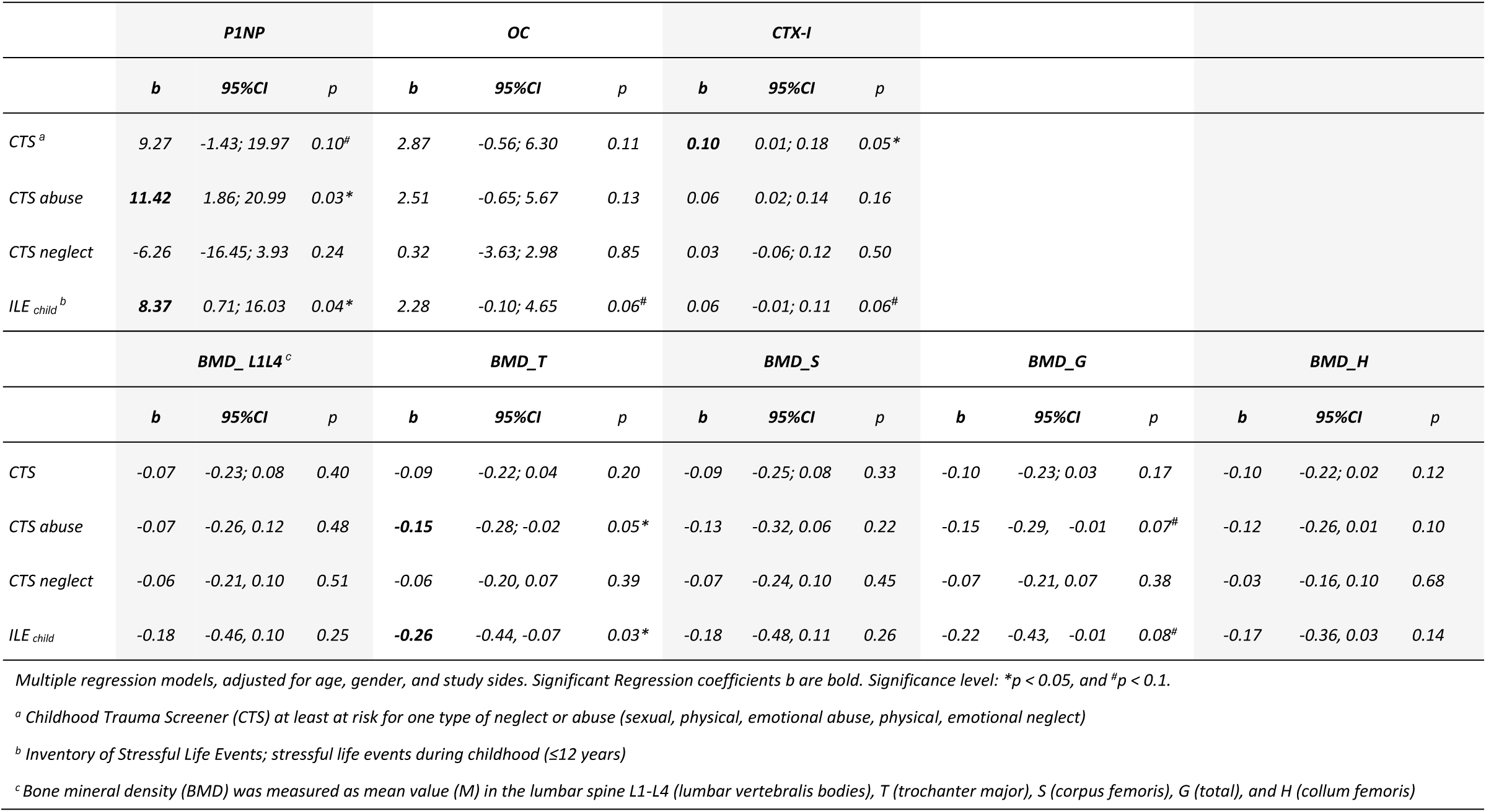
Main effects (regression coefficient b) of childhood maltreatment (abuse or neglect) and stressful life events during childhood (≤12 years; ILE _child_) on bone serum marker (osteocalcin (OC), procollagen type 1 N-terminal propeptide (PINP), and C-terminal telopeptide of type I collagen (CTX-I)) and bone mineral density (BMD)

|  | P1NP |  |  | OC |  |  | CTX-I |  |  |  |  |  |  |  |  |
| --- | --- | --- | --- | --- | --- | --- | --- | --- | --- | --- | --- | --- | --- | --- | --- |
|  | b | 95%CI | p | b | 95%CI | p | b | 95%CI | p |  |  |  |  |  |  |
| CTS<sup>a</sup> | 9.27 | -1.43; 19.97 | 0.10<sup>#</sup> | 2.87 | -0.56; 6.30 | 0.11 | **0.10** | 0.01; 0.18 | 0.05* |  |  |  |  |  |  |
| CTS abuse | **11.42** | 1.86; 20.99 | 0.03* | 2.51 | -0.65; 5.67 | 0.13 | 0.06 | 0.02; 0.14 | 0.16 |  |  |  |  |  |  |
| CTS neglect | -6.26 | -16.45; 3.93 | 0.24 | 0.32 | -3.63; 2.98 | 0.85 | 0.03 | -0.06; 0.12 | 0.50 |  |  |  |  |  |  |
| ILE<sub>child</sub><sup>b</sup> | **8.37** | 0.71; 16.03 | 0.04* | 2.28 | -0.10; 4.65 | 0.06<sup>#</sup> | 0.06 | -0.01; 0.11 | 0.06<sup>#</sup> |  |  |  |  |  |  |
|  | BMD_L1L4<sup>c</sup> |  |  | BMD_T |  |  | BMD_S |  |  | BMD_G |  |  | BMD_H |  |  |
|  | b | 95%CI | p | b | 95%CI | p | b | 95%CI | p | b | 95%CI | p | b | 95%CI | p |
| CTS | -0.07 | -0.23; 0.08 | 0.40 | -0.09 | -0.22; 0.04 | 0.20 | -0.09 | -0.25; 0.08 | 0.33 | -0.10 | -0.23; 0.03 | 0.17 | -0.10 | -0.22; 0.02 | 0.12 |
| CTS abuse | -0.07 | -0.26; 0.12 | 0.48 | **-0.15** | -0.28; -0.02 | 0.05* | -0.13 | -0.32; 0.06 | 0.22 | -0.15 | -0.29; -0.01 | 0.07<sup>#</sup> | -0.12 | -0.26; 0.01 | 0.10 |
| CTS neglect | -0.06 | -0.21; 0.10 | 0.51 | -0.06 | -0.20; 0.07 | 0.39 | -0.07 | -0.24; 0.10 | 0.45 | -0.07 | -0.21; 0.07 | 0.38 | -0.03 | -0.16; 0.10 | 0.68 |
| ILE<sub>child</sub> | -0.18 | -0.46; 0.10 | 0.25 | **-0.26** | -0.44; -0.07 | 0.03* | -0.18 | -0.48; 0.11 | 0.26 | -0.22 | -0.43; -0.01 | 0.08<sup>#</sup> | -0.17 | -0.36; 0.03 | 0.14 |

Multiple regression models, adjusted for age, gender, and study sides. Significant Regression coefficients *b* are bold. Significance level: \**p* < 0.05, and <sup>#</sup>*p* < 0.1.
<sup>a</sup> Childhood Trauma Screener (CTS) at least at risk for one type of neglect or abuse (sexual, physical, emotional abuse, physical, emotional neglect)
<sup>b</sup> Inventory of Stressful Life Events; stressful life events during childhood ( $\leq 12$ years)
<sup>c</sup> Bone mineral density (BMD) was measured as mean value (M) in the lumbar spine L1-L4 (lumbar vertebral bodies), T (trochanter major), S (corpus femoris), G (total), and H (collum femoris)

#### 2.2.3. Bone mineral density

An explorative analysis on bone mineral density in a subsample of DXA measurements suggests that depressive patients *with* stressful life events during childhood (ILE _child_) had reduced BMD in the trochanter major (p = 0.03* for BMD_T) and as a tendency a reduction in general BMD (p = 0.08^#^ for BMD_G, ***Table 2***). There are indications that depressed patients with a history of childhood abuse showed specifically a BMD reduction in the trochanter major (p = 0.05* for BMD_T), and a tendency in reduced BMD in general (p = 0.07^#^ for BMD_G), giving notice of long-term health consequences of people with maltreatment and stressful life events during childhood in comparison to people with depression lacking experiences of early life stress (**Table 2, Figure 5**).

**Figure 5:**
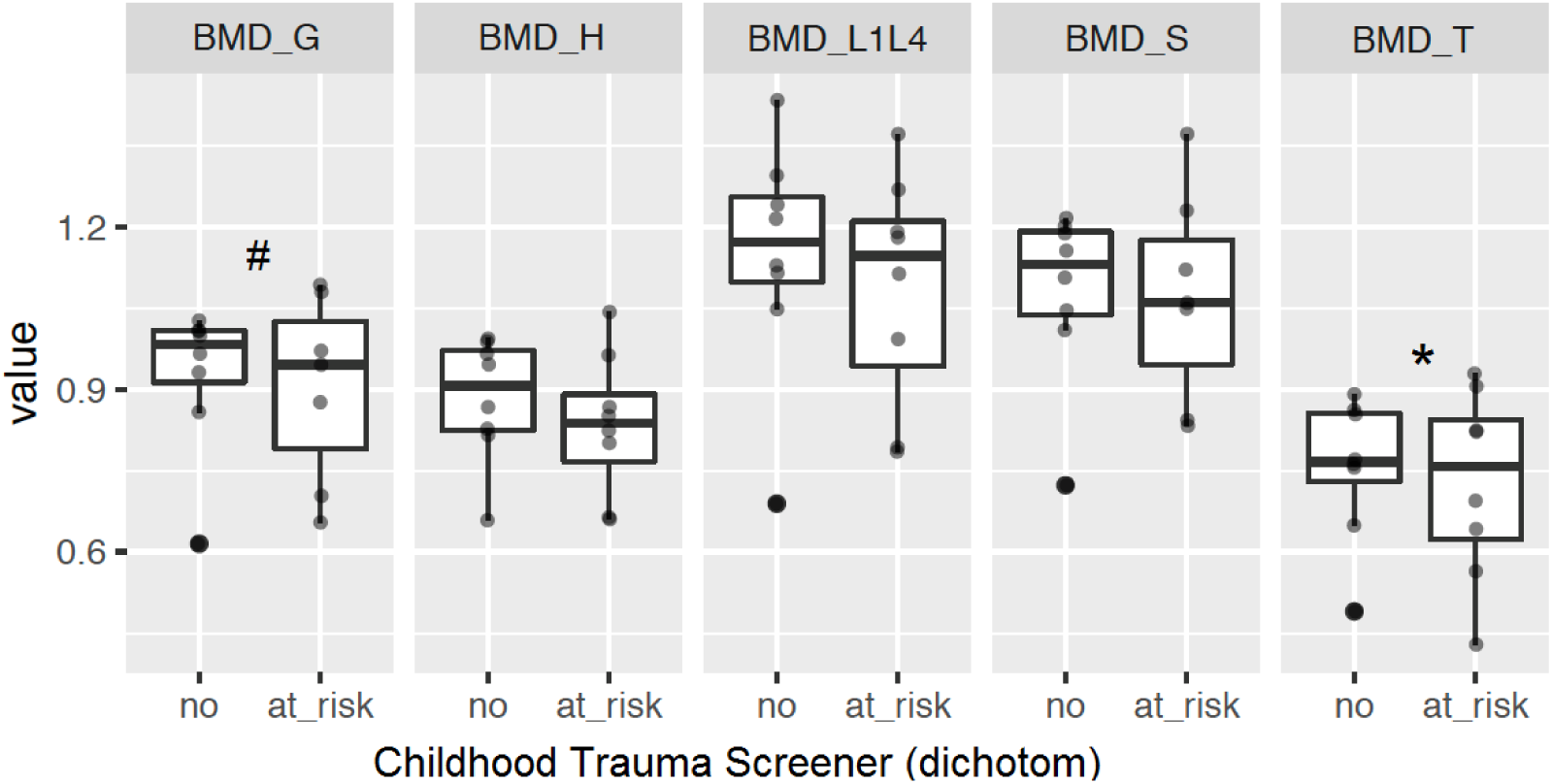
Bone mineral density measurements in human patients. Shown is the BMD in G (total), H (collum femoris), S (corpus femoris), and T (trochanter major) as mean of the right and left site and L1-L4 (lumbar vertebralis bodies) stratified for at risk and not at risk for childhood maltreatment (not controlled for gender, age, body weight). Data (n=18) are expressed as mean and standard deviation of BMD, stratified for CTS (at risk for abuse or neglect). Significance level: *p < 0.05, and ^#^p < 0.1 between indicated groups.

## 3. Discussion

The first aim of this study was to investigate the effects of early life stress on bone health and associated biological factors in a mouse model of early life stress (MSUS). The second study aim was the translation of the gained knowledge in a human model using a sample of depressive patients with and without early life stress.

**Aim 1**: In summary, we found that early life stress in the MSUS model altered bone innervation and neuronal mediator expression as well as bone metabolism, as evidenced by increased CTX-I serum concentrations, but did not affect expression of bone markers within the bone or bone microarchitecture. In detail, we found significantly decreased expression of NGF, NPYR1, VIPR1 and TACR1 in MSUS bones. This response pattern is suggestive for a higher rate of bone turnover and lower bone healing capacities as a consequence of early life stress, suggesting a concurrent joint downregulation, reduced bone remodeling and long-term destabilization. The neurotrophin NGF is important for the expression of cholinergic and sensory neuronal factors and hence for the development and the repair of bone, as demonstrated by reduced bone length when absent [14-26]. The neuropeptide NPY promotes osteoblast differentiation and balances the effects of stress-induced bone-loss through beta-adrenergic stimulation, both with consequences for bone remodeling during healing [10, 53, 56-63]. The neuropeptide VIPR1, finally, promotes bone mineralization [64-68] while TACR1 regulates osteoblasts, osteoclasts and mesenchymal stem cell functionality associated with protection from osteoporosis [2, 62, 69-85].

While MSUS mice showed a clear neuroendocrine response pattern, gene expression of typical bone markers (in bone) was unaffected. Similarly, we did not observe increased osteocalcin in the serum of MSUS mice. However, CTX-I serum levels were significantly increased in MSUS mice, which is an increasingly promoted and robust biochemical factor for bone resorption in clinical settings and further giving notice about correlations to bone mineral density [86-89]. Additional analyses, such as bone histomorphometry to determine whether MSUS leads to alterations in the numbers of osteoblasts (type 1 procollagen staining) and/or osteoclasts (TRAP staining) [90] could provide useful insights into bone adaptation processes.

Regarding bone microarchitecture, no significant changes were observed after normalization to body weight. Body weight itself, which was affected by MSUS, did cause minor changes in bone microarchitecture, likely due to differences in mechanical loading of the skeleton. It is unclear, if altered body weight might be a post stress consequence for MSUS or an incidental finding. This should be considered in future studies, as it can be a confounding factor in micro-CT analyses. To test whether the observed lack of differences in bone microarchitecture (despite an increase in serum CTX-I) might be due to divergent timelines of these events [91] or might be sex-dependent (different susceptibility to bone alterations and/or sensitivity to stress) [92, 93]. We additionally conducted micro-CT scans of the femurs of female mice over a more extended time range (up to 16 months). We did not find any differences in bone microarchitecture in the female mice, even not at later time points (data not shown). It is important to keep in mind that changes in bone microarchitecture may be site specific and may hence have been overlooked by restricting the micro-CT analysis to the femur only. Perhaps, the use of models of aging (e.g., the senescence-accelerated mouse = SAM) [94], models that include bone injury requiring repair (such as osteotomy) [95], or a comparison of stress-resilient and stress-prone subpopulations [96] might potentially have led to observable changes in bone structure within a reasonable time frame [97], but further investigations will be needed to allow for more definite statements.

**Aim 2:** In summary, we found that people that experienced childhood maltreatment or stressful life events during childhood showed an overshooting anabolic bone metabolism during a depressive episode, rather than the expected minor, short-term metabolic adaptation. For example, people in our study showed a threefold higher metabolic reaction in PINP during the depressive episode, in comparison to depressed adults without childhood maltreatment [5]. This was evidenced by increased serum levels in bone metabolism markers (distinctly in PINP and CTX-I, less in OC). Further, our data suggests that depressive adults with experiences of childhood maltreatment (childhood abuse) and stressful life events during childhood showed a significantly decreased BMD in the long term. In contrast, for depressive adults with no experiences of childhood neglect, these patterns were not observed. In other words and regarding latest studies: an overshooting or lacking metabolic adaptation during a depressive episode may serve as a link for the often observed development of BMD in depressed patients in the long term [5]. These mechanisms fit the hypotheses of the destructive effects of a hypersensitive central stress response (overshooting) or a blunted response (lacking adaptation) on organs and bodily tissues.

Other human studies have linked childhood maltreatment to neuroendocrinological and neuroanatomic alterations that overlap with depressive symptoms and result in a vulnerable phenotype that is affected by a hypersensitive central stress response, blunted stress reactivity and lower stress resilience [98-100]. It was suggested that further severe distress, such as additional traumatization, or the development of a depression, may promote an over-adaptive reaction. The response pattern in our data seems to confirm this assumption, which can result from various aspects, such as an altered mitochondrial function, gene dysregulation, accumulation of oxidative stress, inflammation and senescence or changes in neuronal plasticity [27, 28]. The associations between stressful events early in life and an early decrease of noradrenergic, cholinergic and sensory neuronal factors was also observed in our animal model. It is also comprehensible that no such pattern could be found for childhood neglect, as it represents acts of omissions that are typically temporary, such as lacks of nutrition, medical or emotional care. Hence, these might have a lower impact on a person’s psychological integrity, identity and health than childhood abuse. Further, the data matches well with further results within this study sample: patients that suffer from chronic stress (measured by the allostatic load index) showed a catabolic shift in bone metabolism during a depression episode [5]. This can be explained by the fact that other mechanisms are expected for allostatic load as an accumulating damage over time with the consequence of neuroendocrinological and neuroanatomic alterations [36, 99, 101]. The observed reduced bone mineral density can be expected in both response pattern: within a hypersensitive stress response and within an increased allostatic load. Both mechanisms were demonstrated in data of people of the presented DEPREHA-study.

In summary, we found that early life stress affects bone metabolism in mice and humans. In MSUS mice, this was evidenced by a significant shift towards a neurogenic and osteogenic catabolic milieu (decreased NGF, NPYR1, VIPR1, TACR1; higher innervation density in bone; increased serum CTX-I), but no alterations in bone microarchitecture in relatively young mice not exposed to additional life stresses. In humans however, that suffered both from early life stress and depression a significant reduction in BMD was observed. Taking the age range and morbidity (depression) of the human sample under consideration, the differences between mice and human could be interpreted as a patterns of vulnerability and acute consequences (mouse model) compared to long-term burden of early life stress (human). Otherwise, species-related differences in bone metabolism between mice and humans have been described [102]. Our data supports the notion that the mechanisms of bone loss seem different between mice and humans, as mice did not show any decrease in trabecular thickness, which is considered as typical for ageing humans [103]. However, this contrasts Halloran et al. [103], who showed that cancellous bone volume fraction is reduced in 24 month old male C57BL/6J mice and is hence comparable to humans. Furthermore, the observed differences in the response of mice and humans are likely caused by differences in types of stress early in life. In fact, we were able to demonstrate clear differences between different types of early life stress in the human sample, further supporting this theory. Although the results obtained in the mouse study do not overlap entirely with the human results, it provides relevant and promising potential for translation. Furthermore, the animal model can be used to identify probable mechanisms underlying osteoporosis development following early life stress (i.e. related to innervation patterns) that could not be investigated easily in humans, thereby allowing to develop ideas for future studies or even intervention and prevention strategies.

In conclusion, this study demonstrates that stressful life events early in life leads to (mal-) adaptive responses in bone metabolism in both mice and humans. However, the major conclusion to be drawn from the presented data is that different types of early life stress lead to different response patterns concerning bone health and have to be consider separately and possibly consecutively: MSUS mice responded with enhanced bone resorption, which however did not cause changes in bone microarchitecture 8-10 months after the stress exposure when later kept under stress-free conditions. In humans, a reduction in BMD was evident only in depressive patients with severe stress exposure in early life like childhood abuse and stressful life events during childhood. Similarly, the changes in bone serum markers were also dependent on the severity of stress experiences early in life. People with childhood abuse and stressful life events during childhood showed an increased bone metabolism while people with childhood neglect showed a counter regulated reaction. Hence, the consequences of the observed changes in the context of challenges to bone health (fractures, aging etc.) should be studied in a more personalized manner and future prevention strategies should respect the severity of the stress exposure to be treated (e.g. specific type of stress experiences during childhood, double hit paradigms) to achieve the optimal benefit for patients with increased risk for bone pathologies, such as people with depressive disorders.

## 4. Materials and Methods

### 4.1. Mouse Study

#### 4.1.1. Animal model

Animal experiments were conducted in strict adherence to the Swiss Law for Animal Protection and were approved by the local authority (Veterinäramt des Kantons Zürich, Switzerland) under license number 57/2015. C57Bl/6J mice were obtained from Janvier (France) and bred in-house to generate the male mice (n=47 MSUS, n=46 Control) used for experiments.

The MSUS paradigm was conducted as previously described [48]. As shown in ***Figure 6***, 2-3 months old C57Bl/6J naïve females were mated with naïve males for 1 week. Mothers were assigned to control or MSUS group based on the number of male pups born on that day to balance the groups. The MSUS paradigm started on postnatal day 1 (PND1): Pups were separated unpredictably from their mother for 3 hours per day from PND1 until PND14. Separation occurred at any time during the dark cycle. In addition, mothers were subjected to forced swim in cold water (18 °C for 5 min) or restraint in a plastic tube (20 min) at unpredictable times during the 3 hours of separation. Control mice were left undisturbed except for weekly cage changes and weight measurements. Pups were weaned at PND21 and assigned to sex and treatment matched cages (4 to 5 mice) between PND22 and PND28. Co-assignment of siblings was avoided to exclude litter effects. At the age of 8-10 months, control (*n=24*) and MSUS (*n=24*) males were deeply anesthetized by Isoflurane (Attane, Piramal Enteprise Limited), followed by decapitation. Males were single-housed the day before sacrifice to reduce potential acute stress effects [104]. Body weight was monitored throughout the experiment and on the day of single housing. Adult mice were housed in groups of 3 to 5 animals in individually ventilated cages (SealSafe PLUS, Tecniplast). Animals were kept in a temperature- and humidity-controlled facility on a 12h reversed light/dark cycle (light on at 20:00) with food (M/R Haltung Extrudat, Provimi Kliba SA) and water *ad libitum*. Cages were changed weekly.

**Figure 6:**
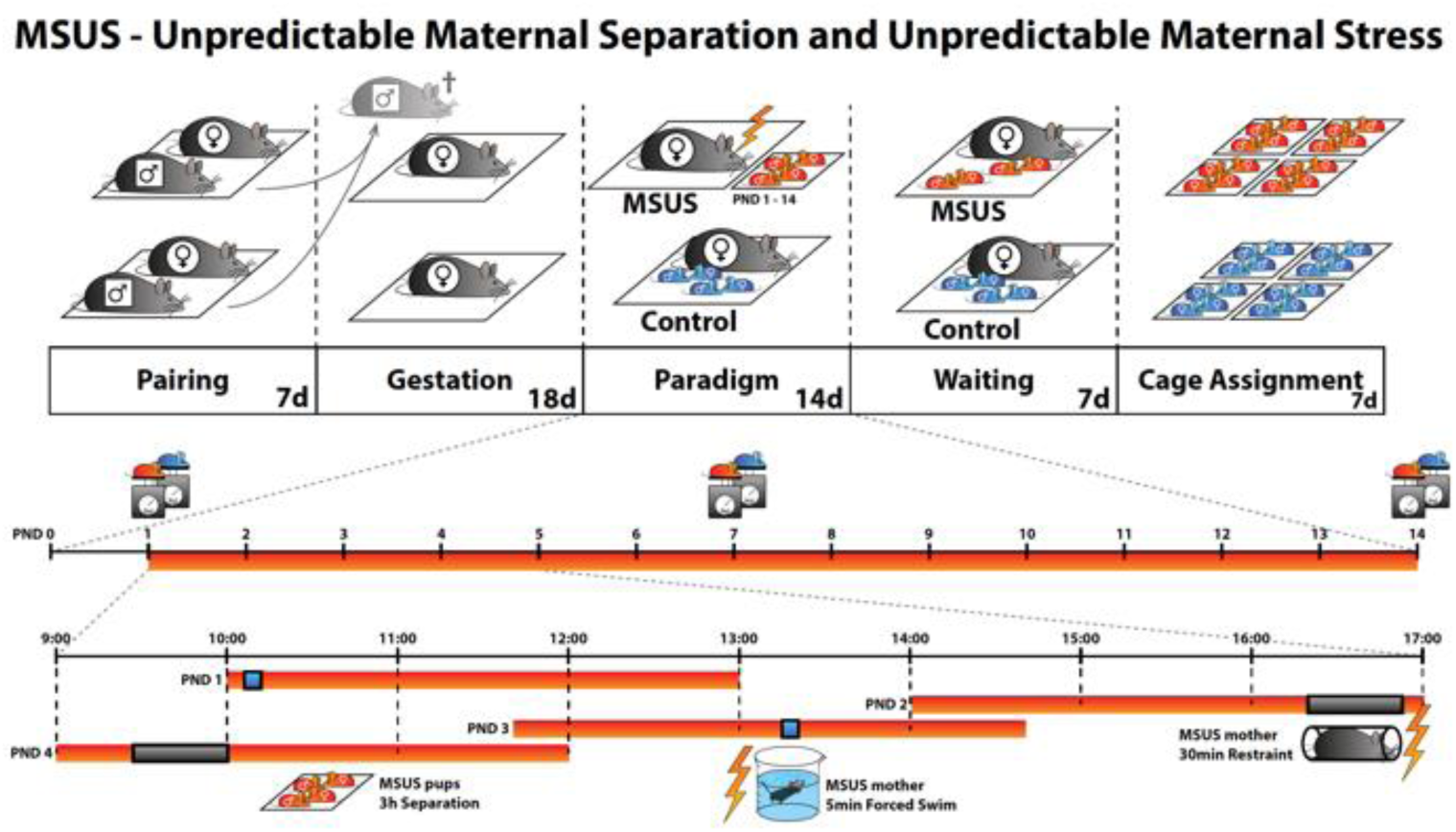
MSUS is a mouse model of early life stress. Unpredictable maternal separation and unpredictable maternal stress is an animal model for early life stress. Naïve males are mated with naïve females for 1-7 days, then males are removed. Dams gestate for about 21 days in normal conditions until delivery. The MSUS paradigm (red bar in the timeline) starts at postnatal day 1 (PND1) and lasts until PND14. During MSUS, pups are unpredictably separated from their mothers for 3 hours each day at different times of the dark cycle (lights off from 8am to 8pm). During separation, the mother is stressed unpredictably by forced swim or restraint as shown in the timeline at the bottom. Pups and dam are left undisturbed from PND15 until weaning at PND21. Gray mice are naïve without any previous stress exposure. Blue pups are controls without any stress exposure. Red pups are exposed to MSUS.

#### 4.1.2. Collection of serum samples

Trunk blood was collected in 2 ml tubes immediately after decapitation and incubated at 4°C. Clotted blood was centrifuged at 2000 rcf at 4°C for 10 minutes. The collected serum was stored at - 80°C.

#### 4.1.3. Collection of bone samples

After euthanasia and blood collection, the left femur was dissected on ice under aseptic conditions and cleaned from muscle tissue. The ends were cut off to allow for removal of bone marrow (flushed out with sterile phosphate buffered saline, PBS) and the entire flushed femur (including the ends) was subsequently shock-frozen in liquid nitrogen and stored at −80°C until further processing for qPCR. Right femurs were dissected, cleaned and stored in 70% ethanol at 4°C until μCT analysis. The third tail bone was dissected and immersed in a mixture of 4% paraformaldehyde and 14% saturated picric acid overnight at 4°C, cryoembedded and stored at −80°C until further processing for histomorphometry was done.

#### 4.1.4. Histomorphometry of neuronal markers

For bone innervation morphometry, immunohistochemistry of pan-neuronal marker and exemplary markers for adrenergic, cholinergic and peptidergic nerve fiber subsets were performed with undecalcified 14 μm thick transverse mouse tail cryostat sections analog to previous descriptions [55]. The sections were done through the center of the third tail bone and contained the bone with the adherent muscle, tendons and skin for internal positive control. Primary antibodies used in this study were: anti-PGP 9.5 (Biotrend, Cologne, Germany), anti-GAP 43 (Millipore, Darmstadt, Germany), anti-ChAT (Serotec, Kidlington, UK), anti-TH (Abcam, Cambridge, MA, USA) and anti-SP (BMA Biomedicals, Augst, Switzerland). Only PGP 9.5 and GAP 43 gave satisfactory staining results, most likely due to insufficient fixation of more digestion prone markers in bone compared to the skin (positive controls were all positive), which commonly require transcardial fixation [105]. Standard controls were performed by omission of the primary antibodies and by incubation with mouse IgG1 instead of primary antibodies. No labeling was observed in negative controls. Nerve fiber profiles were counted per area with the help of a grid equipped ocular at x200 magnification using a Leica inverted automated fluorescence-microscope (Leica, Wetzlar, Germany). Briefly, numbers of positively stained fibers were counted per grid field in at least 30 consecutive fields per experimental mouse and in at least *n=8* mice per experimental group by two blinded independent researchers. The data were then pooled and expressed as means per group +/- standard error of the mean.

#### 4.1.5. qPCR for gene expression in bone samples

Mouse femurs were pulverized using a metal ball mill (Retsch, Germany) and total RNA was isolated with Trizol/chloroform extraction, followed by column-based purification using RNeasy Mini Kit (Qiagen, Germany) according to the manufacturer’s protocol, but including proteinase K incubation (55°C, 15 min). Contaminating DNA was removed by using the RNAse-free DNAse kit (Qiagen, Germany) and 32 μl of RNA with an average yield of 48 ng/μl were reverse transcribed with Superscript II reverse transcriptase (Life Technologies, Germany) to obtain 80 μl of cDNA. qPCR analysis was performed in duplicate, using the TaqMan Fast PCR Master Mix with Taqman assays for bone markers (Life Technologies, Germany) (***Supplementary Table 2***) or the QuantiTect PCR Kit with self-designed primers for neuronal factors (Qiagen, Germany) (***Supplementary Table 3***). The amount of cDNA per reaction depended on the respective target gene and its expression level and was 1-18 ng. Gene expression was normalized to the housekeeping gene (dCt) and 2^-dCt^ were computed.

#### 4.1.6. ELISA of serum samples

Commercially available ELISA kits were used to determine the serum levels of OC (60-1305, Immunotopics/Teca, Switzerland), PINP (AC-33F1, Immunodiagnosticsystem, Germany) and CTX-I (AC-06F1, Immunodiagnosticsystem, Germany) according to the manufacturers’ instructions. Serum was used either undiluted (CTX-I) or at a 1/11 (osteocalcin) or 1/10 (PINP) dilution.

#### 4.1.6. Micro-CT of bone samples

Analysis of bone microarchitecture by μCT was conducted as previously described [106]. Briefly, bones were measured on a μCT40 (Scanco Medical AG, Brüttisellen, Switzerland), operated at 55 kVp and 145 μA with an integration time of 200 ms and 2-fold frame averaging. Images were reconstructed at an isotropic nominal resolution of 10 μm and filtered using a constrained 3D Gaussian filter (sigma 0.8, support 1) to suppress noise. Masks of the full bone, cortical (shaft at 56% bone length, 1 mm long) and trabecular region of interest (from 66% to 88% of bone length) were created automatically. Morphometric parameters were calculated according to standard guidelines such as total volume (TV), bone volume (BV), marrow volume (MV), apparent volume density (AVD), femur length, cortical area fraction (Ct.Ar/Tt.Ar), cortical thickness (Ct.Th), trabecular bone volume fraction (BV/TV), specific bone surface (BS/BV), trabecular thickness (Tb.Th), trabecular spacing (Tb.Sp) and trabecular number (Tb.N) [107]. Due to differences in body weight between the control mice and the MSUS mice, data was normalized for body weight and body weight adjusted means were compared.

### 4.2. Human Study

#### 4.2.1. Participants

*n*=240 patients with depressive disorder (ICD-10 F32.x, F33.x, F34.1, F43.21) were recruited of which *n*=208 completed the initial examination (baseline (t_0_), for more details see [5]. A subsample of *n*=54 patients took part at the fourth follow up measurement (t_4_) including DXA imaging. Patients who fulfilled the following criteria were included: 18 to 65 years of age, depressive episode (ICD-10 F32.x or F33.x), dysthymia (F34.1) or an adjustment disorder with prolonged depressive reaction (F43.21), ≥ 21 days absenteeism within the last year. Exclusion criteria were: pregnancy, hormonotherapy (expect contraceptive and thyroid hormone therapy), inability to fill in a questionnaire, intellectual disabilities (ICD-10 F70-79) or one of the following diseases: acute infection, endocrine and metabolic disorders, neurological diseases, dementia (ICD-10 F00-F03), schizophrenia (ICD-10 F20), emotional-unstable personality disorders (ICD-10 F60.3x), disease of the immune system, substance abuse and dependency (expect nicotine).

All participants were fully informed in verbal and written form about the intent and content of the study, and gave their written informed consent. The clinical investigations were conducted according to the principles of the Declaration of Helsinki. Final ethical approval was provided on 11/12/2017 from the Ethics Review Board of the University of Potsdam, Germany (number 15/2017).

#### 4.2.2. Study Procedure

Study objectives were investigated at two assessments (t_0_ and t_4_) of an 8-month observational multicenter study with four measurement points: baseline (t_0_), after 5 weeks (t_1_), 5 months (t_2_), 8 months (t_3_) and an additional follow-up measurement (t_4_: M= 15.4 month, SD = 4.5 month after baseline). At each measurement point, study participants answered a comprehensive questionnaire (t_0_ – t_4_) under the supervision of trained study nurses. The questionnaire comprised demographic characteristics, psychological symptoms (e.g. depression) and constructs (e.g. stress, early life events, affect, coping), physical ailments, and information regarding alcohol consumption and medication intake. Blood samples were collected at baseline (t_0_) and 5 months follow-up (t_2_) and DXA bone densitometry measurements took place at the last follow up (t_4_).

#### 4.2.3. Psychometric Measures

Early life stress was measured retrospectively comprising experiences of childhood maltreatment and stressful life events during childhood. The Childhood Trauma Screener (CTS) [108], a 5-item screening tool of the 28-item Childhood Trauma Questionnaire (CTQ) [109, 110], assesses five types of childhood maltreatment such as emotional, physical, and sexual abuse plus emotional and physical neglect. Each item is scored on a five-point rating scale from “never true” (1) to “very often true” (5). In accordance to Glaesmer et al. [111], we classified participants at risk if they rated at least mild forms of childhood abuse or neglect. We additionally controlled for response bias by the 3-item Minimization-Denial subscale from the CTQ and excluded participants when indicated. Cronbach’s Alpha was previously specified with 0.76 [112].

Stressful life events were assessed by a modified version of the Inventory of Stressful Life Events (ILE; [113]). Participants rated 34 adverse life events regarding occurrence, frequency and year of occurrence. The accumulation of stressful, life-changing events at different stages of development was counted for every participant, in this analysis focusing on childhood (ILE _child_ ≤ 12 years). The scores ranged between 0 (no strain) and 4 (high strain). Cronbach’s Alpha for ILE was 0.83. Depressive symptoms and severity were assessed using the Beck Depression Inventory-II (BDI-II) [114, 115]. Internal consistency in the sample was Cronbach’s Alpha 0.89.

Furthermore, confounding factors like sociodemographic and biometric characteristics and study site were assessed.

#### 4.2.4. Serum bone marker measurement

Blood samples were drawn from the arm in the morning (7-9 am), collected in plain blood collection tubes, allowed to clot at room temperature for 30 min, followed by centrifugation, isolation of serum and freeze-storage until further analysis. Bone-related blood markers (OC, PINP, CTX-I) were analyzed in serum samples by electrochemiluminescence immunoassays “ECLIA” (12149133 122 for Osteocalcin, 03141071 190 for PINP, 11972308 122 for CTX-I, all from F. Hoffmann-La Roche, Ltd., Basel, Switzerland)

#### 4.2.5. DXA measurement

Bone mineral density (BMD) was measured by DXA bone densitometry measurement (Lunar, Prodigy Advance, GE Healthcare, Illinois, USA) in the lumbar spine (lumbar vertebral bodies L1-L4) and both hips. Parameters are given as mean values (M) of the left and right site: G (total), H (collum femoris), S (corpus femoris), T (trochanter major), and L1L4 (lumbar vertebral bodies L1-L4).

### 4.3. Statistical Analysis

Data consistency was checked, screened for outliers and analyzed descriptively. Continuous variables were also tested for normality by using skewness, kurtosis, omnibus test; variance homogeneity was proven by Variance Ratio and Levene test.

For the examination of **Aim 1** (mouse model), student t-test for equal or Aspin-Welch Unequal Variance test were performed, using multiple t-test and adjusted for multiple comparison by Holm-Sidak method (alpha = 0.05). Since body weight was found to be significantly different, ANCOVA models with body weight as covariate were applied for micro-CT data to compute corresponding means at their covariate means (i.e. least square means at covariate mean for body weight of 16.41). In addition, blocking on total body weight was used as a means of accounting for collinearity in analyses of variance and mediation analyses were conducted using linear regression models to identify mediation effects between body weight and bone microstructure. All reported tests are two-sided, and p-values < 0.05 are considered as statistically significant.

Regarding **Aim 2** (human model) multiple regression models were applied. Statistical models were cross-sectional and controlled for age, gender, study sites and additionally body weight for DXA measurements.

Statistical analyses were conducted using NCSS (NCSS 10, NCSS, LLC. Kaysville, UT), STATISTICA 13, IBM SPSS 25 and the statistical software R [116, 117].

## Supplementary Materials

Supplementary materials can be found at www.mdpi.com/xxx/s1.

## Author Contributions

KW-K planned and coordinated the project, participated in tissue collection, supervised experiments and drafted the manuscript; MR conducted the animal experiments, participated in tissue collection and drafted the manuscript; EC participated in tissue collection, analyzed samples and drafted the manuscript; AB helped to coordinate the human study, analyzed sample and proof-read the manuscript; GAK participated in tissue collection, analyzed samples and drafted the manuscript; TA analyzed samples and proof-read the manuscript; WH conducted statistical analysis of all animal experiments and proof-read the manuscript; DD conducted statistical analysis of all human experiments and proof-read the manuscript; RM provided input in micro-CT analysis and proof-read the manuscript; MAR helped in the coordination of the human study and proof-read the manuscript; IMM planned and coordinated the animal study, supervised experiments, acquired funding and proof-read the manuscript; EMJP helped to coordinate the mouse study, analyzed samples and drafted the manuscript; PMW planned and coordinated the project, conducted human study as PI, supervised experiments, acquired funding and drafted the manuscript.

## Acknowledgments

In this section you can acknowledge any support given which is not covered by the author contribution or funding sections. This may include administrative and technical support, or donations in kind (e.g., materials used for experiments).

## Abbreviations

ADRB2: adrenoceptor beta 2
ANCOVA: analysis of covariance
AVD: apparent volume density
BDI-II: Beck Depression Inventory Revision
BDNF: brain derived neurotrophic factor
BMD: bone mineral density
BMD_G: bone mineral density general
BMD_H: bone mineral density collum femoris
BMD_L1L4: bone mineral density lumbar vertebralis bodies 1-4
BMD_S: bone mineral density corpus femoris
BMD_T: bone mineral density trochanter major
BV: bone volume
BS: specific bone surface
CGRP: calcitonin gene related peptide
ChAT: choline Acetyltransferase
CHRNA7: cholinergic Receptor Nicotinic Alpha 7 Subunit
CTQ: Childhood Trauma Questionnaire
CTS: Childhood Trauma Screener
CTX-I: c-terminal telopeptide of type I collagen
Ct.Ar: cortical area fraction
Ct.Th: cortical thickness
DAPI: 4′,6-diamidino-2-phenylindole
dCt: delta Ct
DEPREHA: [Der Einfluss unterschiedlicher Behandlungs-settings auf den Therapieerfolg bei Patienten mit depressiven Erkrankungen in der Rehabilitation]
DXA: Dual Energy X-ray Absorptiometry
ECLIA: electrochemiluminescence immunoassays
ECM: extracellular matrix
ELISA: enzyme-linked immunosorbent assay
FITC: Fluorescein isothiocyanate
GAP 43: Growth Associated Protein 43
HPA: hypothalamic pituitary adrenal
ICD-10: International Classification of Diseases Version 10
IgG1: Immunoglobulin G
ILE: Inventory of Stressful Life Events
ILE child: Inventory of Stressful Life Events; life events during childhood ≤12 years
μCT: micro-computed tomography
M: mean
MV: marrow volume
MSUS: maternal separation and unpredictable stress
mRNA: messenger ribonucleic acid
NF: nerve fibers
NGF: nerve growth factor
NPY: neuropeptide Y
NPYR1: neuropeptide Y receptor 1
OC: osteocalcin
qPCR: quantitative polymerase chain reaction
PBS: phosphate buffered saline
PINP: procollagen type 1 N-terminal propeptide
PND: postnatal day
RAMP1: receptor activity modifying protein 1
SAM: senescence-accelerated mouse
TAC1: tachykinin 1
TACR1: tachykinin receptor 1
Tb.N: trabecular number
Tb.Sp: trabecular spacing
Tb.Th: trabecular thickness
TH: tyrosine hydroxylase
TRKA: tropomyosin receptor kinase A
Tt.Ar: total area
TRKB: tropomyosin receptor kinase B
TV: total volume
VIP: vasoactive intestinal peptide
VIPR1: vasoactive intestinal peptide receptor 1

## Supplementary Data

### Supplementary Figures

**S-Figure 1:**
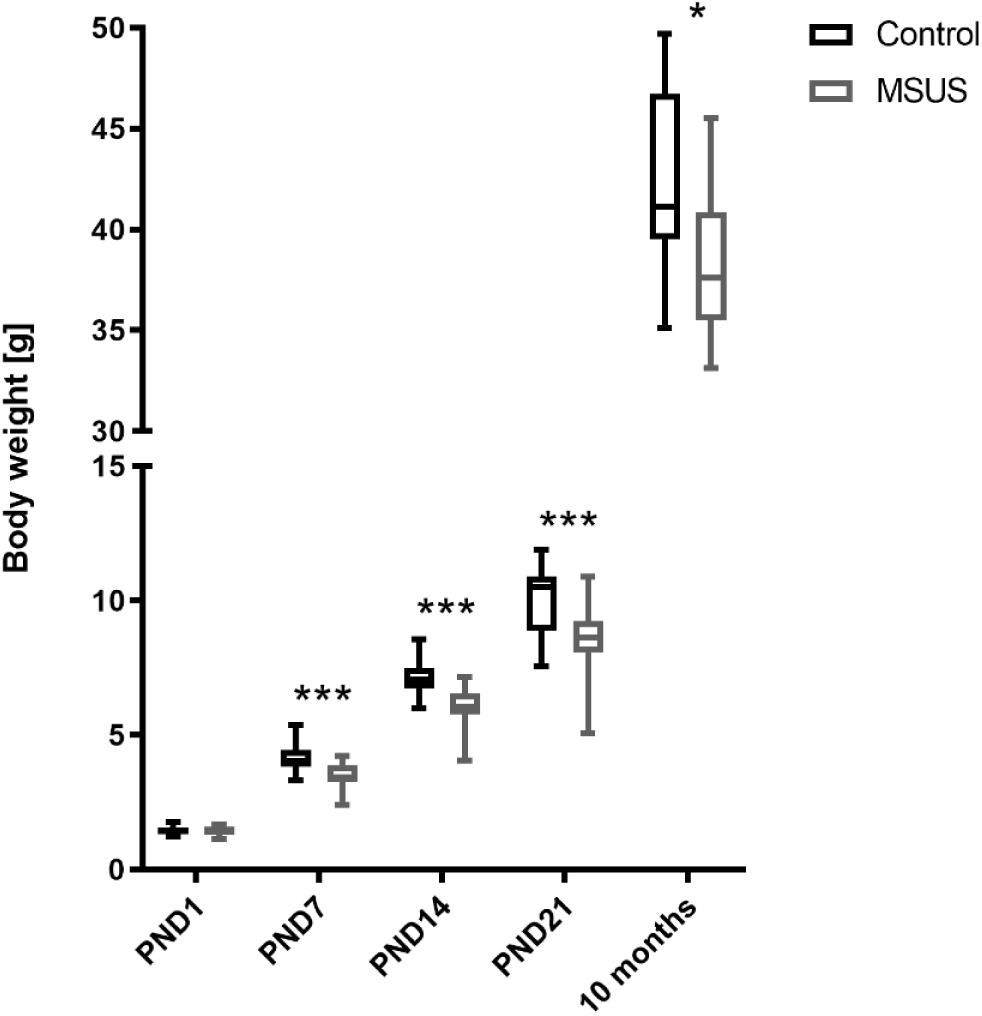
Body weight development. Shown are body weights of male pups, which are equal at PND1 and decreased following MSUS at PND7 (n=45 Control, n=46 MSUS), PND14 (n=45 Control, n=46 MSUS), PND21 (n=46 Control, n=44 MSUS) and in adults (n=14 Control, n=14 MSUS). Data are expressed as Min to Max of body weight in [g] and considered significant at *p < 0.05 and ***p < 0.001.

### Supplementary Tables

**S-Table 1:**
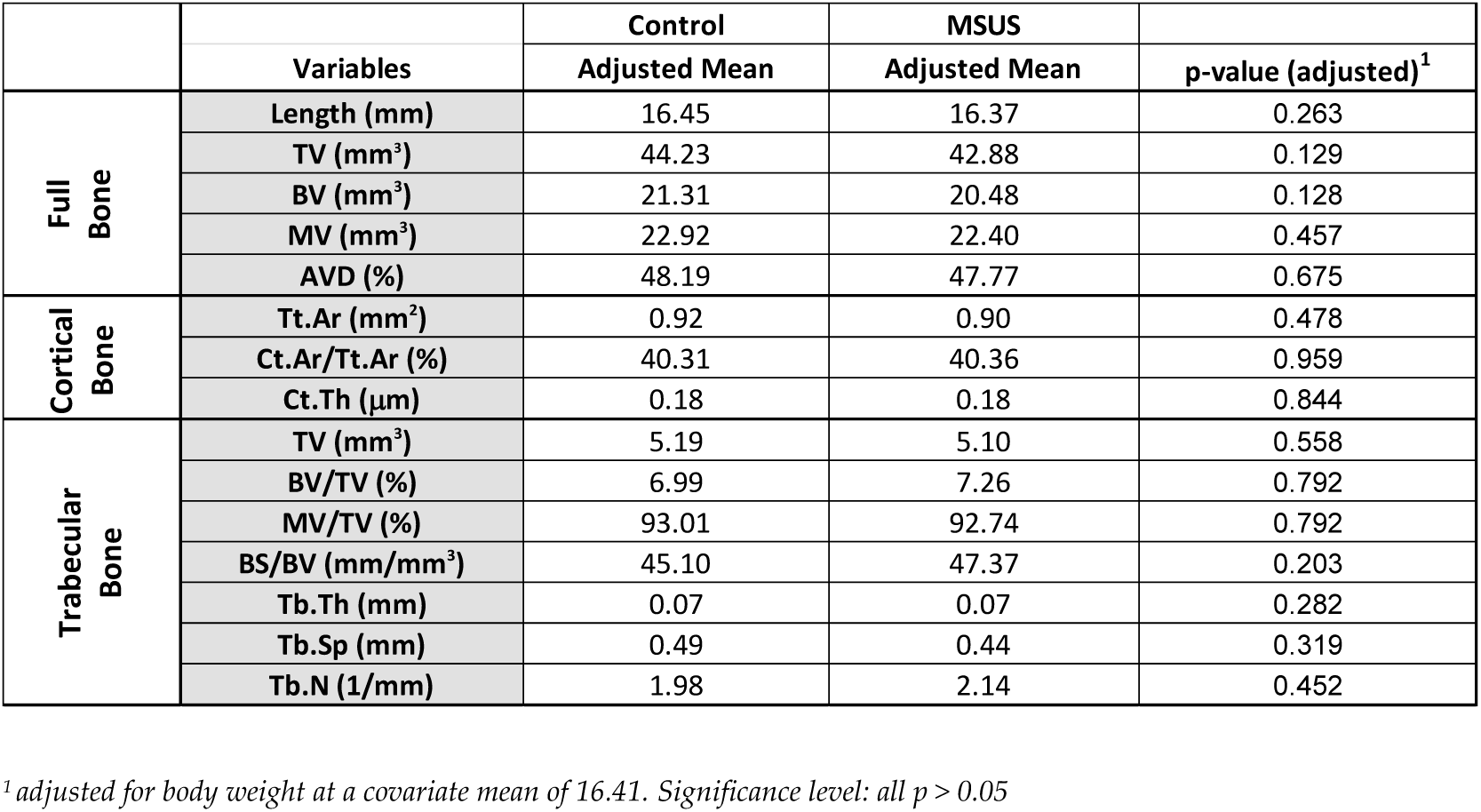
Micro-CT data obtained for full, cortical (Ct) and trabecular (Tb) bone in control mice and MSUS mice with regard to length, total volume (TV), bone volume (BV), marrow volume (MV), apparent volume density (AVD), area (Ar), thickness (Th), bone surface (BS), spacing (Sp) and number (N) and thereof calculated characteristics.

|  |  | Control | MSUS |  |
| --- | --- | --- | --- | --- |
|  | Variables | Adjusted Mean | Adjusted Mean | p-value (adjusted)<sup>1</sup> |
| Full Bone | Length (mm) | 16.45 | 16.37 | 0.263 |
|  | TV (mm<sup>3</sup>) | 44.23 | 42.88 | 0.129 |
|  | BV (mm<sup>3</sup>) | 21.31 | 20.48 | 0.128 |
|  | MV (mm<sup>3</sup>) | 22.92 | 22.40 | 0.457 |
|  | AVD (%) | 48.19 | 47.77 | 0.675 |
| Cortical Bone | Tt.Ar (mm<sup>2</sup>) | 0.92 | 0.90 | 0.478 |
|  | Ct.Ar/Tt.Ar (%) | 40.31 | 40.36 | 0.959 |
|  | Ct.Th (μm) | 0.18 | 0.18 | 0.844 |
| Trabecular Bone | TV (mm<sup>3</sup>) | 5.19 | 5.10 | 0.558 |
|  | BV/TV (%) | 6.99 | 7.26 | 0.792 |
|  | MV/TV (%) | 93.01 | 92.74 | 0.792 |
|  | BS/BV (mm/mm<sup>3</sup>) | 45.10 | 47.37 | 0.203 |
|  | Tb.Th (mm) | 0.07 | 0.07 | 0.282 |
|  | Tb.Sp (mm) | 0.49 | 0.44 | 0.319 |
|  | Tb.N (1/mm) | 1.98 | 2.14 | 0.452 |

<sup>1</sup> adjusted for body weight at a covariate mean of 16.41. Significance level: all $p > 0.05$

**S-Table 2:** Taqman gene expression assays.

| Target | Identifier |
| --- | --- |
| GAPDH | Mm99999915_g1 |
| Osteocalcin | Mm03413826_mH |
| Osteoprotegerin | Mm00435452_m1 |
| Osteopontin | Mm00436767_m1 |
| Sclerostin | Mm00470479_m1 |

**S-Table 3:**
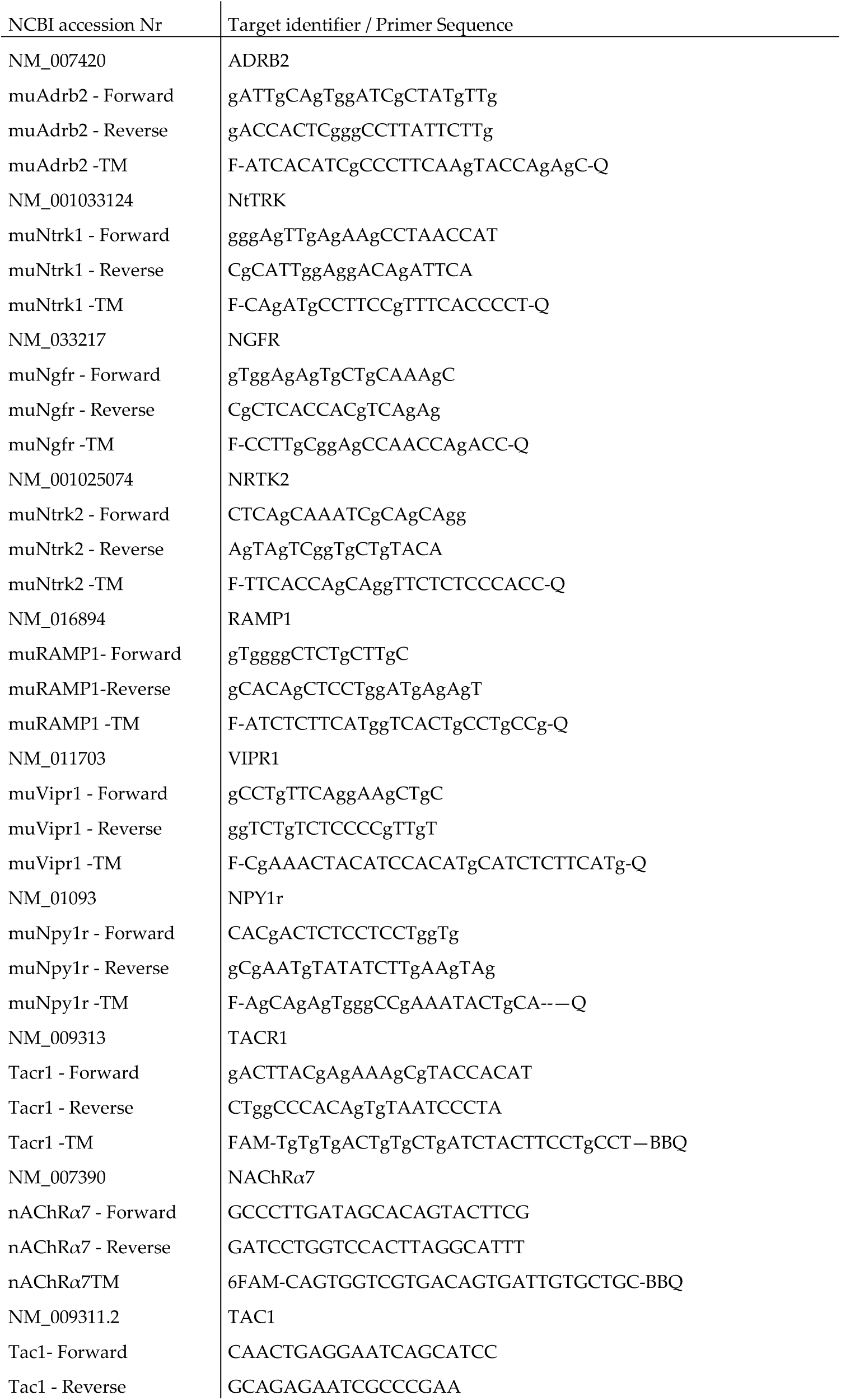
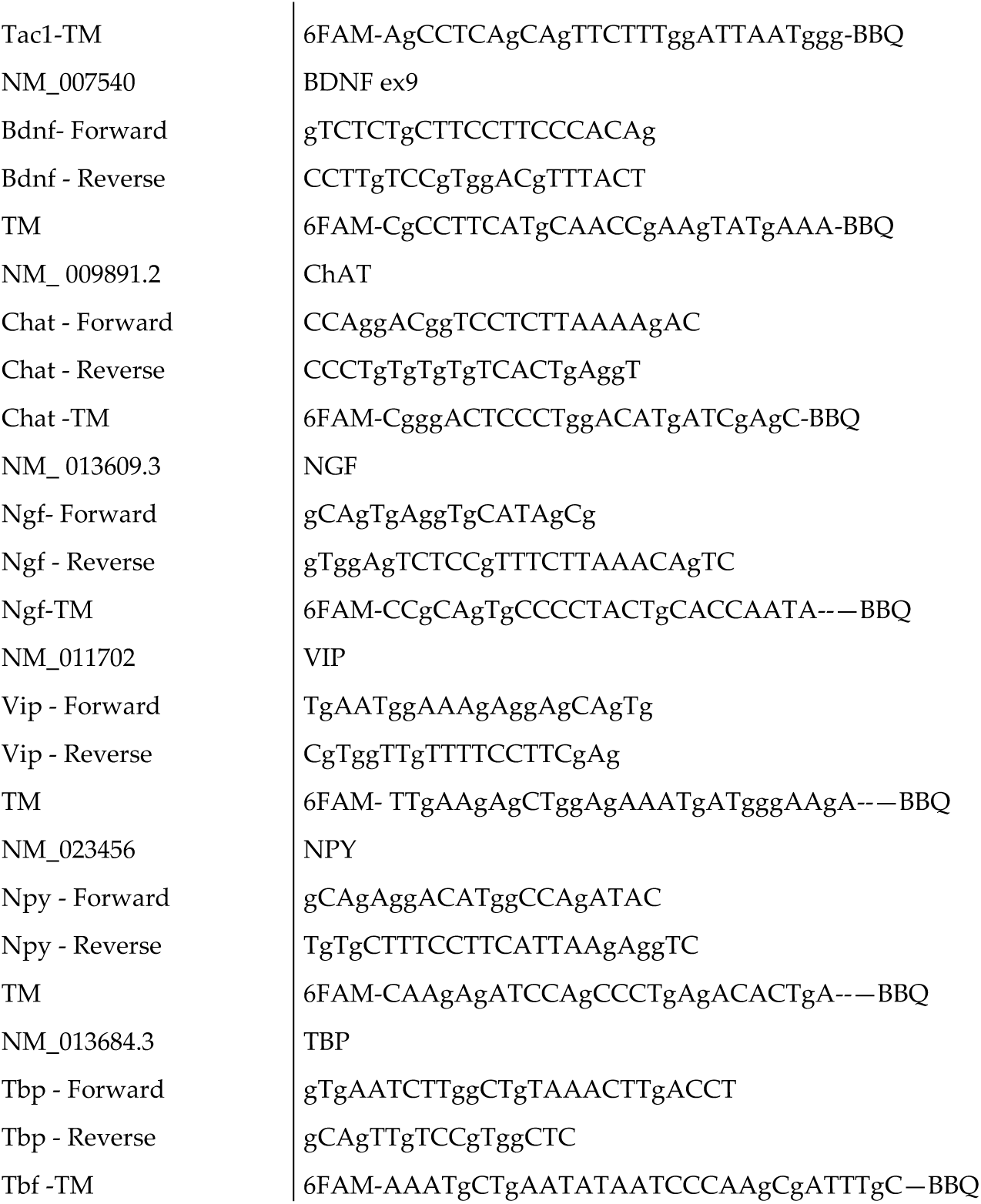
Self-designed primers.

